# A method to screen for meiotic drive using embryonic markers

**DOI:** 10.64898/2026.05.22.727232

**Authors:** Benjamin K. McCormick, Daniel A. Barbash

## Abstract

Meiotic drivers are selfish elements which co-opt gametogenesis to increase their own transmission. Driving alleles may spread in a population even if harmful to overall fitness, requiring the emergence of host suppressors to ameliorate these costs. In some cases, intense co-evolutionary arms races between drivers and their host genomes may occur. These conflicts have been invoked to explain the rapid evolution of karyotypes and reproduction-associated proteins across the tree of life. Despite their evolutionary importance, relatively few meiotic drivers have been well-characterized, in large part due to the difficulties inherent to detecting meiotic drivers, distinguishing them from viability effects, and performing systematic screens. To address these gaps, we present an approach to driver detection at the embryo stage in wild-derived *D. melanogaster*. By combining fluorescent markers with a method to induce embryonic arrest at a standard developmental stage, we detect transmission of wild-derived alleles as compared to their fluorescently marked homologs in early embryos, before most fitness differences among alleles (which may mimic drive) manifest. We provide proof-of-concept for the approach and identify several areas for future improvement.

## Introduction

Meiotic drivers are selfish elements that bias the process of gametogenesis so that they are overrepresented among functional gametes (*e.g.* Courret *et al*. 2019; Searle and Pardo-Manuel 2024; Presgraves *et al*. 2026). This property allows driving alleles to spread in a population even if they harm their hosts, leading to arms race dynamics between drivers and the genomes that harbor them. Meiotic drive conflicts are believed to have important implications for genome evolution, from reshaping the karyotype (Blackmon *et al*. 2018) to driving accelerated evolution of proteins involved in conserved processes like reproduction (Chang *et al*. 2023) and chromosome segregation (Roach *et al*. 2013). Despite their evolutionary significance, the diversity of mechanisms by which drivers act, their genomic signatures, and their frequencies and patterns of variation in natural populations remain almost completely unknown in even the best studied models. This is in large part because characterized drivers, while wide in their phylogenetic distribution, are quite uncommon. It is, however, unclear if this is reflective of actual scarcity or simply arises from biases in driver detection.

Historically, most meiotic drive alleles have been discovered fortuitously, usually through linkage of a strong distorter to either a sex chromosome or visible marker (Courret *et al*. 2019). It thus stands to reason that the pool of known drivers is likely highly biased in multiple respects, potentially obscuring important patterns of driver mechanism and evolution. While systematic screens are lacking, several methods have been proposed for the detection of new meiotic drivers, though none have been widely applied. First, pooled sequencing of adult offspring from haplotype-resolved F1s derived from inbred parental lines can be used to screen genome-wide for evidence of drive (Wei *et al*. 2017). However, this approach is resource-intensive and thus limited in throughput. Additionally, biases and noise arising from sources both technical (e.g. sequencing biases) and biological (e.g. differential viability among genotypes) may mimic or obscure true drive. Furthermore, many drivers are linked to lethal alleles or are themselves lethal in homozygotes (Larracuente & Presgraves 2012; Presgraves *et al*. 2026), introducing inherent bias to approaches which can only be applied to inbred lines. Another general approach to driver detection is to screen for signatures associated with known mechanisms of drive and suppression. For example, a proposed bioinformatic signature of male meiotic drive is the existence of tandem arrays of homologous genes on opposing sex chromosomes (Ellison & Bachtrog 2019), especially when one array is made up of protein-coding genes and the other gives rise to testis-expressed short RNAs (e.g. siRNAs or piRNAs) against these genes.

Similarly, one could also screen for cytological phenotypes associated with drive, like failure of some proportion of spermatids to undergo individualization, as occurs for many known male drivers in *Drosophila*. However, such approaches, while useful, are limited to discovery of systems with familiar architecture and mechanisms of drive and suppression. To address these and other limitations of current methods, we present and provide proof-of-concept for a straightforward approach to conducting a (mostly) unbiased genetic screen of chromosomes derived from outbred or wild-derived *Drosophila melanogaster* for evidence of transmission distortion.

## Results and Discussion

### Using centromere-linked markers to screen for meiotic drive

To overcome the limitations of existing methods for meiotic driver discovery, we sought an approach that is: 1) sensitive and high throughput; 2) unbiased with respect to driver mechanism; 3) amenable to use with outbred wildtype flies; 4) works on frozen samples without any cryopreservation; 5) usable for screening either male or female meiotic drive; and 6) able to distinguish drive from most causes of differential viability. For our initial tests, we generated a stock carrying an easily-scorable dominant fluorescent marker in tight linkage to the chromosome 2 centromere (Fig. 1). This marker stock can be crossed to wild-derived lines to generate transgene-heterozygous F1s to be screened for drive. The transmission ratio of the marker in the F2 generation reflects the inheritance of the marked centromere region in females or the entire chromosome in males (due to the lack of meiotic recombination in males) relative to its wild-derived homolog (Fig. 1). Thus, if a driver is linked to the wildtype copy of the chromosome 2 centromere, this will manifest as an under-recovery of the fluorescent marker in F2 flies. Note that the scheme depicted in Fig. 1 assumes that the starting wild stock is inbred. If the wild-derived stock to be screened is instead outbred, one would likely want to first isolate a single chromosome over a balancer to ensure that only one isolate of a chromosome is screened at a time.

**Figure 1:**
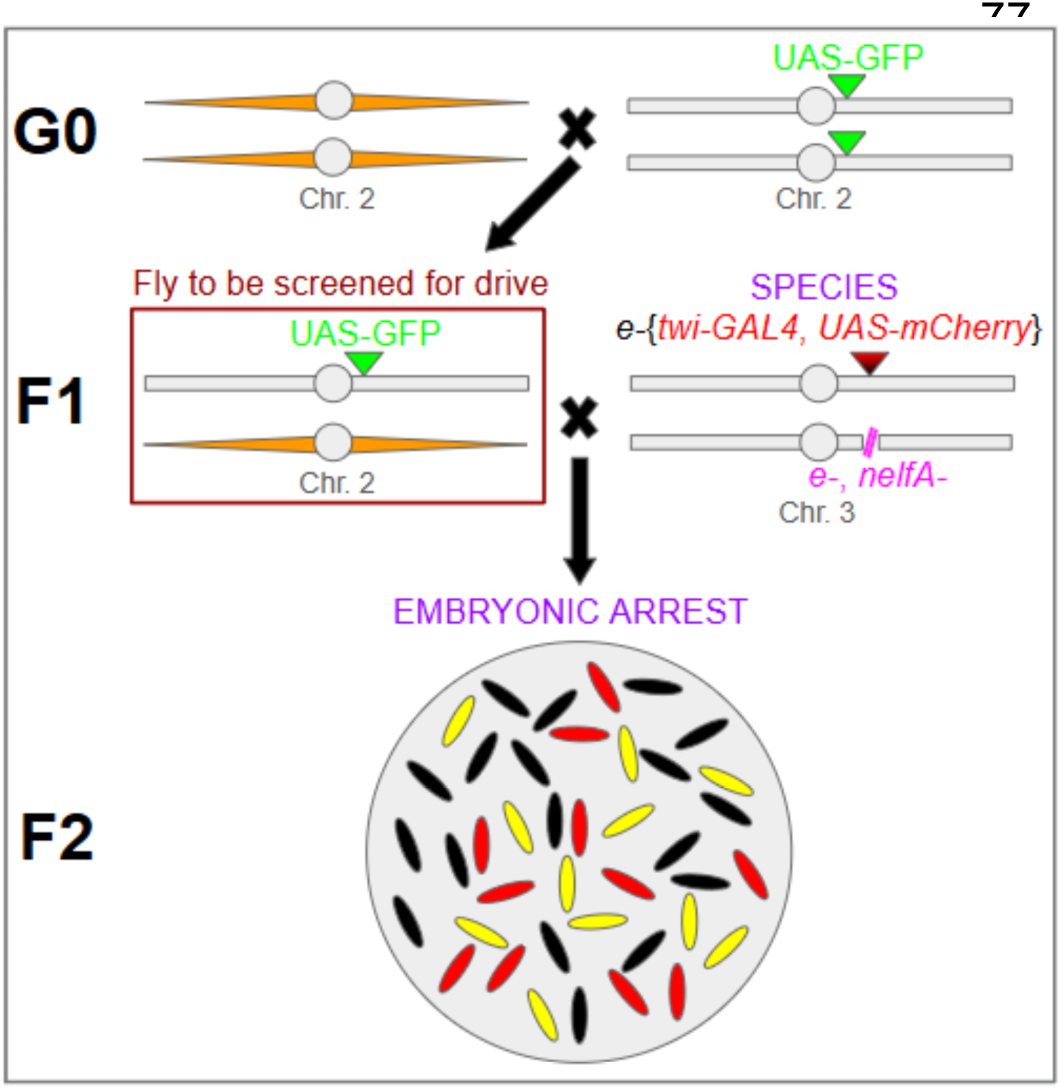
Crossing scheme to screen a single copy of the chromosome 2 centromere for evidence of meiotic drive. In the first generation, a stock carrying the chromosome to be screened, shown in orange, is crossed to a homozygous marker stock carrying the *UAS-tdGFP* transgene. The resultant F1s are then crossed to SPECIES lines carrying the *twi-GAL4, UAS-mCherry* transgene (note that this allele is balanced over an overlapping homozygous lethal deficiency, shown in pink). F2 embryos resulting from this cross will die during early development due to the dominant lethality produced by outcrossing the SPECIES stocks. Embryos can then be mounted on slides, imaged, and scored for red and green fluorescence (see Figure 2). In the absence of drive, among RFP+ embryos (i.e. those fertilized embryos which inherited the *twi-GAL4, UAS-mCherry* transgene), equal numbers of GFP^+^ RFP^+^ (yellow) and GFP^−^ RFP^+^ (red) embryos are expected. On the other hand, if the chromosome screened produces meiotic drive, an excess of GFP^−^ RFP^+^ embryos will be observed. Unfertilized eggs and the 50% of embryos that do not inherit the *twi-GAL4, UAS-mCherry* transgene will lack fluorescence (black). Note that the same approach could be applied to any locus in the genome, depending on the insertion site of the *UAS-tdGFP* marker.

Even in the absence of meiotic drive, non-Mendelian transmission ratios can be caused by differences in viability among alleles. Distinguishing drive from viability effects is best done by examining offspring allele frequencies early in development, during embryogenesis. Embryonic screening of *Drosophila*, however, presents several challenges due to their rapid developmental rate. Chief among these is the need to standardize the developmental timing of scoring to ensure that embryos are not scored earlier or later than the marker is expressed, which would result in systematic undercounting of marker-containing embryos, falsely mimicking drive. To accomplish this without the need for tightly timed embryo collections, which would substantially reduce the throughput of the assay, we sought to induce embryonic arrest at a consistent developmental stage. We made use of a set of stocks called Synthetic Postzygotic barriers Exploiting CRISPR-based Incompatibilities for Engineering Species (SPECIES; Buchmann *et al*. 2021). These are stable lines that, when outcrossed to any other strain of *D. melanogaster*, result in inviable offspring that die as embryos. This dominant lethality is caused by the SPECIES lines expressing a nuclease-deficient deactivated Cas9 (dCas9) protein fused to a transactivation domain as well as several gRNAs that target the regulatory regions of essential developmental genes. The SPECIES lines themselves carry protective indels interrupting the target sites of these gRNAs, but when they are outcrossed to wildtype animals lacking these indels, the resulting strong overexpression of the targeted genes in early development of the progeny causes embryonic lethality. In our scheme, we outcross the marker heterozygotes to be screened for drive to a modified SPECIES strain, thus inducing developmental arrest in the embryos to be scored.

Other requirements of using embryos are that the fluorescent markers 1) exhibit strong expression in heterozygous early embryos to allow easy and reliable scoring, ideally without the need for dechorionation and 2) lack maternal expression, since otherwise all embryos from heterozygous mothers would exhibit fluorescence regardless of zygotic genotype. We tested fluorescent transgenes driven by a variety of regulatory constructs and found that *twist (twi)-GAL4* combined with *UAS*-driven fluorescent reporters meets these requirements. We therefore marked the chromosome 2 centromere with *UAS-tdGFP* and integrated a *twi-GAL4* transgene in the SPECIES line to which the marker heterozygotes are crossed (Fig. 1, S1).

An additional complication is that *D. melanogaster* females often lay unfertilized eggs which, if counted as non-fluorescent embryos, would give a false signal of drive. To address this problem, we modified the *twi-GAL4* transgene in the SPECIES stock to also contain *UAS-mCherry*. All progeny embryos from a SPECIES parent inheriting the *twi-GAL4, UAS-mCherry* transgene will fluoresce red while unfertilized eggs remain non-fluorescent. We then measure the ratio of GFP+:RFP+ embryos which is expected to be 1:2 in the absence of drive regardless of whether or not unfertilized eggs are present (see Fig. 2).

**Figure 2:**
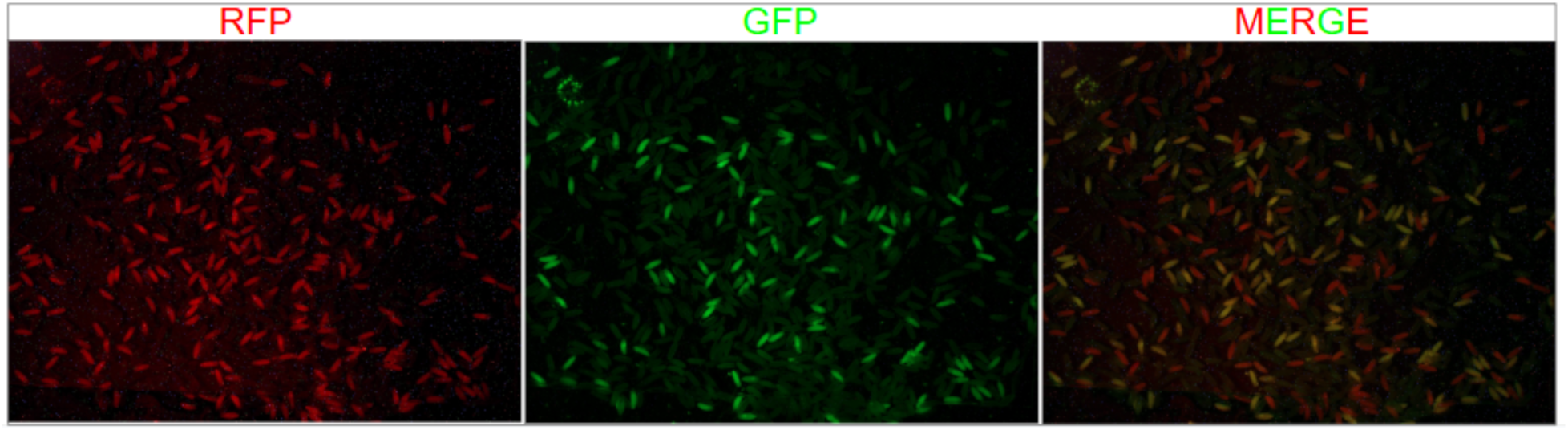
Representative images of a slide of embryos from a cross of B04/2R1 *UAS-tdGFP* F1 females crossed to SPECIES *twi-GAL4, UAS-mCherry*/+ males. No drive occurs in this cross. Panels from left to right: red fluorescent channel, green fluorescent channel, merged image. Camera settings used are in Supplemental File 3.

The combined *twi-GAL4, UAS-mCherry* transgene transformed into the SPECIES stock is homozygous lethal for unknown reasons, which created two challenges. First, the transgene must be balanced in some manner to be maintained as a stock. Because the SPECIES lines cannot be outcrossed, an existing balancer chromosome cannot be introduced. To circumvent this problem, we engineered via CRISPR in the SPECIES strain a homozygous-lethal deletion overlapping the site where the *twi-GAL4, UAS-mCherry* transgene is integrated (Fig. S1A) and can thus maintain both the transgene and the deletion in a stable transheterozygous state (Fig. S1B). Second, since the transgene remains heterozygous in the SPECIES line, it will only be inherited by half of the progeny, and only these progeny can be scored.

As a proof of concept, we applied our assay using the chromosome 2 marker to two inbred wildtype stocks from the Global Diversity Lines (GDL; Grenier *et al*. 2015) as well as a line carrying *Segregation Distorter* (*SD*), a known male meiotic driver on chromosome 2 (Larracuente & Presgraves 2012), which we used as a positive control (Table 1). We observed Mendelian segregation of all chromosomes tested relative to the marker regardless of the cross direction, with the exception of males carrying the *SD* chromosome where, as expected, inheritance was significantly nonrandom (72% transmission of *SD*; p = 0.0008). Progeny numbers in male drive crosses were, however, very low. This is because after several years of maintenance of the *twi-GAL4, UAS-mCherry* transgene in the SPECIES background we found 1) significantly reduced female fecundity (the primary cause of low sample sizes) and 2) partial breakdown of arrest, with occasional offspring hatching as larvae (1.7-7.0%; Supplemental File 1, Table 1) and rarely even surviving to adulthood. Extensive tests of the stock when first constructed suffered from neither issue. These issues may have arisen from off-target effects of the dCAS9/gRNAs responsible for producing embryonic arrest and/or compensatory changes to the genome to cope with such effects.

**Table 1:** Genotype counts from crosses of males and female F1 marker heterozygotes to track the inheritance of each chromosome screened (the chromosomes tested were derived from two Global Diversity lines, I17 and B04; SD[72], a known male drive allele on chromosome 2, was used as a positive control. Recovery of the unmarked autosome screened for drive was calculated for each cross by dividing the count for the RFP+, GFP- class by the total number of RFP+ offspring observed. To test for evidence of drive, we performed a 2×2 Chi-squared test on the observed counts (GFP+ and GFP-) among RFP+ embryos compared to Mendelian expectation (1:1 recovery of both products). Additional underlying data including counts of hatched larvae can be found in Supplemental File 1, Table 1.

| Sex | Source of chromo-<br>some<br>screened | RFP+,<br>GFP-<br>(source<br>chr. 2) | RFP+,<br>GFP+<br>(marked<br>chr. 2) | Total<br>RFP+ | Total<br>embryos<br>and<br>larvae | Expected<br>transmis-<br>sion of<br>source<br>chr. 2 | Observed<br>transmis-<br>sion of<br>source<br>chr. 2 | p-value |
| --- | --- | --- | --- | --- | --- | --- | --- | --- |
| Female | B04 | 553 | 531 | 1084 | 2324 | 0.5 | 0.47 | 0.62 |
| Male | B04 | 26 | 23 | 49 | 107 | 0.5 | 0.53 | 0.69 |
| Female | I17 | 398 | 402 | 800 | 1689 | 0.5 | 0.50 | 0.92 |
| Male | I17 | 18 | 20 | 38 | 92 | 0.5 | 0.47 | 0.82 |
| Female | SD[72] | 271 | 282 | 553 | 1161 | 0.5 | 0.49 | 0.74 |
| Male | SD[72] | 84 | 33 | 117 | 370 | >>0.5 | 0.72 | 0.00080 |

We attempted to address the reduced female fecundity by outcrossing to the parental SPEICES line (B2), intercrossing the F1 progeny, then selecting for transgene-deletion transheterozygotes in the F2 generation. We then tested females from this rebuilt stock against males carrying a chromosome two extracted from an isofemale line collected in Zambia (Lack *et al*. 2015). From three plates of collection we observed very few hatched larvae and obtained 3251 total eggs/embryos, 1330 of which were Rfp^+^. Among the Rfp^+^ embryos, 669 were Gfp^+^, for an observed recovery of 0.503 (p = 0.89). These results demonstrate that we were able to restore robust female fertility to the SPECIES line containing *twi-GAL4, UAS-mCherry* and obtain a high rate of arrest. Together with the data in Table 1, our results highlight the sensitivity and accuracy of the approach and underscore the robustness of the fluorescent markers.

### Limitations and Future Directions

Meiotic drive is typically inferred from progeny ratios in F1 adults but is caused by events that occurred many days prior during meiosis or gametogenesis in their male or female parent. This leaves a high chance of conflating meiotic drive with post-zygotic viability effects that differ between alleles. We developed here a method to screen for drive during embryogenesis, which greatly reduces the potential for artifacts caused by viability differences. Our method uses genetic arrest of all developing embryos and is not affected by potential differences in developmental rate among genotypes, because embryos can be allowed to continue development for at least 24 hours prior to harvesting. Our method also allows easy detection of fluorescent markers from either fresh or frozen embryos without the need for dechorionation or any other processing.

Embryonic scoring in Drosophila poses a particular problem in that females often lay unfertilized eggs that cannot be easily distinguished from arrested embryos without detailed examination. For meiotic drive screening, undetected unfertilized eggs would be mistakenly classified as Gfp-negative embryos and give an inflated signal of drive. We therefore used *twi-GAL4* to express *UAS-mCherry* in all developing embryos such that unfertilized eggs remain non-mCherry. We included both components on a single transgene to ensure they are always co-inherited. Unexpectedly, the transgene is homozygous lethal. It is unlikely that the insertion is directly causing lethality because it is integrated into an *attP* site disrupting the *ebony* gene that is itself homozygous viable. Furthermore, an identical *twi-GAL4* transgene lacking *UAS-mCherry* is also viable in homozygotes when integrated at the *attP40* landing site. Therefore, we speculate that the high dosage of mCherry being expressed during early development is causing lethality. A possible solution would be to attempt to reduce its expression by reducing the number of *UAS* repeats in the *UAS-mCherry* construct.

A caveat to our approach for distinguishing unfertilized eggs is that any embryos that begin development but arrest and die before *twi* expression starts at the blastoderm stage (Jiang *et al*. 1991) will be erroneously classified as being unfertilized eggs. If significant early embryonic lethality were associated with the wild (non-Gfp) chromosome being tested, it would be incorrectly interpreted as negative drive (under-transmission).

During the course of our experiments, we observed reduced fertility and arrest efficiency of our transformed SPECIES stock. We were able to restore the stock by outcrossing the transgene into the original untransformed SPECIES stock, but it does suggest the possibility that the embryonic arrest will lose efficacy over time. Possible alternatives to induce arrest might be to use maternally expressed GAL4 to drive UAS-RNAi constructs targeting essential zygotic embryonic genes (Staller *et al*. 2013), or compound autosome stocks that produce lethal haploid and triploid embryos when crossed to normal euploid stocks (Merrill *et al*. 1998).

Despite some limitations, the system we developed is readily expandable. UAS-Gfp transgenes could be easily placed on other chromosomes and locations. Automation of scoring using either traditional image segmentation or AI workflows would also considerably improve screening efficiency. Using other fluorescent reporters that emit in non-overlapping wavelengths would allow two regions or chromosomes to be screened simultaneously. The ability to detect two different transgenes would also facilitate using a similar embryonic arrest system to quantitate rates of rearrangements or recombination in large numbers of embryos.

## Materials and Methods

PCRs were carried out using iProof™ High-Fidelity DNA Polymerase (Bio-Rad), gel purification steps were carried out using Monarch™ DNA Gel Extraction Kit (NEB), gateway cloning reactions were carried out using Gateway™ LR Clonase™ II Enzyme mix, and ligation reactions were carried out using T4 DNA ligase (NEB) according to manufacturer instructions. The sequence of all plasmids injected was confirmed by Plasmidsaurus sequencing (Supplemental File 1, Table 5). Proper integration of transgenes in the *D. melanogaster* genome was confirmed via PCR as described below. Sequences of primers and additional information about their use can be found in Supplemental File 1, Table 4. CRISPR gRNAs and homology arm sequences can be found in Supplemental File 1, Table 2.

### Fly crosses

Flies were maintained and crossed at 25°C. Because all stocks to be screened (Supplemental File 1, Table 1) were either sufficiently inbred (GDL lines) or already maintained over a balancer (*SD[72]*/*CyO*), there was no need to extract a single copy of chromosome 2 prior to starting the cross scheme (Fig. 1). Briefly, virgin females of each of these strains were mated to chromosome 2 centromere *UAS-tdGFP* marker stock males. *UAS-tdGFP*/+ F1s of both sexes were separately collected and mated to SPECIES B2 *e-{3xP3-dsRed, twi-GAL4, 5xUAS-mCherry-NLS}/e-,Nelf-A-{3xP3-tdGFP}* of the opposite sex in groups of 25 3-5 day old males and females. Embryos were collected in 24 hour windows on grape agar plates with yeast paste, aged for an additional 24 hours, then frozen at -80°C until scoring. Crosses were maintained in this manner for 10 days before being discarded. Embryos were subsequently removed from plates, washed with PBST, and mounted on slides in PBST in groups of up to 1000. We captured fluorescent green and red images of each slide and counted embryos and larvae from these images manually (see Supplemental File 1, Tables 1 and 3 for raw count data and associated camera settings, respectively).

### Construction of CRISPR plasmid for integrating *attP*, *3xP3-dsRed* in the *ebony* CDS

We constructed a plasmid for integration of an *attP* site marked with *3xP3-dsRed* via CRISPR targeting the coding sequences of the gene *ebony* (*e*). The gRNA used was obtained from Kane *et al*. (2017). Two dsDNA constructs were synthesized: *attL3-tRNA-gRNA-Left Homology Arm-attL4(rev)* and *attL5-Right Homology Arm-attL6(rev)* (Twist Bioscience; Supplemental File 1, Table 2) and cloned into the vector *p{dU6-3-GW1-Amp-ccdB-attP-B3-3xP3-dsRed-B3-GW2-Chlor-SacB}* (gift from D. Stern; also known as pJAT7 (Addgene #204292)) in a single step via double gateway cloning as described by Stern et al. (2023). The final vector contains both a gRNA expression system as well as an *attP-3xP3-dsRed* HDR donor construct flanked by appropriate homology arms. It will be referred to as *p{dU6-3-ebony gRNA1-e LHA-attP-B3-3xP3-dsRed-B3-e RHA}* (Supplemental File 1, Table 5).

### Construction of pDEST_APIGH_twi-GAL4_5xUAS-mCherry-NLS

A 3.2 kb fragment of the *twist* (*twi*) promoter approximately corresponding to the 3.1 kb fragment described in Greig & Akam (1993) was amplified from *D. melanogaster* genomic DNA (primers: 2159 + 2160; restriction sites included in 5’ overhangs). This PCR product was then gel purified and ligated into the KpnI and XbaI sites of *pDEST_APIGH_cIAB5(-)*, a plasmid containing an *attB* site, *miniwhite*, and a *gypsy insulator-IAB5 enhancer-hsp70 promoter-GAL4-gypsy insulator* construct. The resulting plasmid, referred to as *pDEST_APIGH_twi-GAL4*, contains the insert ligated between one of the gypsy insulators and *hsp70min-GAL4*, replacing the *IAB5* enhancer sequence. Next, a 1.6 kb fragment containing *5xUAS-hsp70-mCherry-SV40 NLS-SV40 Poly(A) signal* was amplified from *pUAS-mCherry-NLS* (addgene #87695; primers: 2209 + 2210; restriction sites included in 5’ overhangs). *pDEST_APIGH_twi-GAL4* was digested with EcoRV, liberating most of the miniwhite transgene; a 13.2 kb fragment of the backbone was retained and used for cloning the *5xUAS-mCherry* insert. The finished plasmid is referred to as *pDEST_APIGH_twi-GAL4_5xUAS-mCherry-NLS* (Supplemental File 1, Table 5).

### Generation of SPECIES lines carrying *twi-GAL4_5xUAS-mCherry-NLS* constructs

*p{dU6-3-ebony gRNA1-e LHA-attP-B3-3xP3-dsRed-B3-e RHA}* was subsequently injected into SPECIES B2 embryos alongside *Pnanocas9* (11H10; Rainbow Transgenic Flies Inc.). Screening for transformants and subsequent generation of a homozygous stock was accomplished by mating G0s back to SPECIES B2, then sibling mating the F1 offspring in pools of 10. After mating, the F1 flies were PCR screened (also in pools of 10; primers: 2247 + 2230) for presence of the properly integrated transgene. Positive crosses were retained and resulting *e+* F2s were mated in pairs and individually PCR screened. Crosses with two positive (i.e. transgene heterozygous) parents were retained. F3s exhibiting the phenotypes associated with homozygous disruption of *e* were mated in pairs to establish a homozygous stock. Several F4 flies from each cross were spot checked with primers flanking the CRISPR cut site (primers: 2241 + 2242) to ensure that all offspring were homozygous for the intended edit. This stock will be referred to as SPECIES B2 e-{3xP3-dsRed, attP} (27_12).

Subsequently, *pDEST_APIGH_twi-GAL4_5xUAS-mCherry-NLS* was injected into SPECIES B2 e-{attP} alongside the phiC31 integrase expression vector *pBS130* (Rainbow Transgenic Flies Inc.). G0s were mated to one another in groups of four. Vials were checked for embryos and early-stage larvae exhibiting red fluorescence. These embryos/larvae were collected, and once mature, mated in pairs to SPECIES B2. Fluorescent F2 siblings collected as embryos were then mated in pairs. No homozygous F3s were obtained across multiple replicates, indicating that the transgene is homozygous lethal, likely due to strong overexpression of the fluorescent reporter in the early embryo. As such, to balance this allele, we generated via CRISPR an independent deletion (marked with 3xP3-tdGFP) of *e* as well as the neighboring essential gene, *Nelf-A* (Fig. S1).

To accomplish this, we constructed two CRISPR vectors, each expressing a different gRNA, with the guides targeting unique sites 8.9 kb apart in the *e* and *Nelf-A* coding sequences. These vectors were otherwise identical to one another, each carrying a *3xP3-tdGFP* marker flanked by homology arms matching the sequences up- and downstream of the intended deletion (Supplemental File 1, Tables 2 and 5; Fig. S1A). To construct these vectors, we first amplified a fragment containing *3xP3-tdGFP* flanked by SacI and EcoRV sites from *pHD-Gypsy_AttP-Dfd-3xP3-tdGFP* by PCR (primers: 2292 + 2293), which was then ligated into *p{dU6-3-GW1-Amp-ccdB-attP-B3-3xP3-dsRed-B3-GW2-Chlor-SacB}* between SacI and StuI sites, replacing *3xP3-dsRed* with *3xP3-tdGFP*. The resulting plasmid is referred to as *p{dU6-3-GW1-Amp-ccdB-3xP3-tdGFP-GW2-Chlor-SacB}*. Next, gRNA and homology arms constructs were synthesized (Twist Biosciences) and recombined into *p{dU6-3-GW1-Amp-ccdB-3xP3-tdGFP-GW2-Chlor-SacB}* via double gateway cloning, as in the construction of *p{dU6-3-ebony gRNA1-e LHA-attP-B3-3xP3-dsRed-B3-e RHA}* described above (Supplemental File 1, Table 2). The resulting plasmids are referred to as *p{dU6-3-ebony gRNA2-e/Nelf-A LHA-3xP3-tdGFP-B3-e/Nelf-A RHA}* and *p{dU6-3-Nelf-A gRNA-e/Nelf-A LHA-3xP3-tdGFP-B3-e/Nelf-A RHA}* (Supplemental File 1, Table 5). These plasmids were coinjected with *Pnanocas9* (11H10; Rainbow Transgenic Flies Inc.) into SPECIES B2. Next, we selected GFP+ F1 offspring and confirmed the presence of the intended deletion via PCR spanning both expected transgene-genome junctions (primers: 1736 + 2375 for the *Nelf-A* junction; and 1255 + 2247 for the *e* junction). A single double-PCR positive F1 was then mated back to SPECIES B2, and the resulting F2 offspring crossed to the heterozygous *twi-GAL4_5xUAS-mCherry-NLS*/*+* . (F2 offspring were also separately mated to one another to confirm the expected homozygous lethality of the deletion through lack of phenotypically *e-* F3s.) As both the *e-{3xP3-dsRed*, *twi-GAL4*, *5xUAS-mCherry-NLS}* and the *e-,Nelf-A-{3xP3-tdGFP}* alleles generated are homozygous lethal, we were able to construct a stable heteroallelic stock, which will be referred to as *SPECIES B2 e-{3xP3-dsRed, twi-GAL4, 5xUAS-mCherry-NLS}/e-, Nelf-A-{3xP3-tdGFP}* (Fig. S1B).

### Construction of CRISPR plasmid for integrating *UAS-tdGFP* in the pericentric heterochromatin of chromosome 2

A 0.4 kb fragment containing *5xUAS-hsp70* was amplified from pUAS-mCherry-NLS (primers: 2213 + 2214; restriction sites included in 5’ overhangs) and gel purified. The vector pUC57-2R1_Sqh_GFP contains a *gypsy insulator-Sqh promoter_5’ UTR-tdGFP-Sqh 3’ UTR-gypsy insulator* construct flanked by homology arms targeting a locus in the pericentromeric heterochromatin of the right arm of chromosome 2. This vector was digested with PstI and NaeI, while the PCR-generated insert was digested with NsiI (which shares a common 3’ overhang with PstI), leaving one sticky end and one blunt end. Note that the blunt end contains an intact AflII site to be used in the next step). Both of these digests were then gel purified, with a 6.2 kb fragment of the vector backbone containing everything but the *Sqh* promoter and 5’ UTR retained. The insert was then directionally ligated into this vector and cloned. This vector is referred to as pUC57-2R1_intermediate.

A 1.8 kb fragment containing *tdGFP-SV40 Poly(A) signal* was then amplified from pHD_attP_Dfd_Gypsy_tdGFP (primers: 2217 + 2218; restriction sites included in 5’ overhangs). This insert was digested with AflII and BglII, while the vector was digested with AflII and BamHI. Both of these digests were then gel purified, with a 4.5 kb fragment of the vector containing everything but *tdGFP* and the *Sqh 3’ UTR* retained. The insert was then directionally ligated into pUC57-2R1_intermediate between the AflII site (introduced in the last cloning step) and a BamHI site such that, between the two homology arms, the final construct is as follows: *gypsy insulator-5xUAS-hsp70-tdGFP-SV40 Poly(A) signal-gypsy insulator*. This vector will be referred to as pUC57-2R1_UAS-tdGFP (Supplemental File 1, Table 5).

### Making *UAS-tdGFP* chromosome 2 marker stock

pUC57-2R1-UAS-tdGFP was injected into embryos of nos-Cas9 attP2 alongside pCFD3_2R1 (Supplemental File 1, Table 5), a plasmid that expresses the 2R1 gRNA under control of a U6 promoter (Rainbow Transgenic Flies Inc.). G0s were mated to flies carrying a *P{tubP-GAL4}LL7* transgene inserted on the third chromosome (BDSC #5138). F1 larvae were then screened for green fluorescence; positive larvae were allowed to mature then mated to ywF10 (1_43). Nonfluorescent yellow-bodied (i.e. lacking *P{tubP-GAL4}LL7* and y+ from Cas9 transgene) F2s were then mated to a chromosome 2 balancer stock with a dominant visible marker (1_38; yw; CyO balancer). Once having mated, the F2 parents were PCR screened with a primer set with one primer within and one primer outside of the insert (2223 + 2215) to ensure proper integration of the transgene. For positive F2 crosses, F3 flies exhibiting the balancer phenotype were sibling mated in pairs. After mating, the F3 parents were individually PCR screened (2223 + 2215); crosses with 2 positive parents were retained. Next, F4 flies lacking the balancer phenotype were sibling mated in pairs to establish a homozygous stock. After mating, these flies were screened with primers flanking the CRISPR cut site (2201 + 2202) to ensure that both parents were homozygous for the intended edit. To construct an inbred version of this line, we crossed marker males to Canton S females, then sibling mated for 10 generations, selecting visually for recovery of wildtype *white* (*w*) and *yellow* (*y*) alleles and again selecting for homozygosity of the marker via PCR. The resultant stock will be referred to as 2R1 UAS-tdGFP.

## Data availability

All data required to reproduce the analyses and figures presented here can be found in the supplemental files.

## Acknowledgments

We thank Dr. Omar Akbari for sending us the SPECIES stocks, Dr. David Stern for the kind gift of unpublished plasmids, and Dr. John Pool for the Zambia isofemale line. We also thank Ms. Shuqing Ji for construction of the *Nelf-A-, e-{3xP3-tdGFP}* allele as well as collection of the Zambia stock drive data and Mr. Ethan Lai for early tests of the suitability of various embryo-expressed markers for drive screening.

---

**Figure S1:**
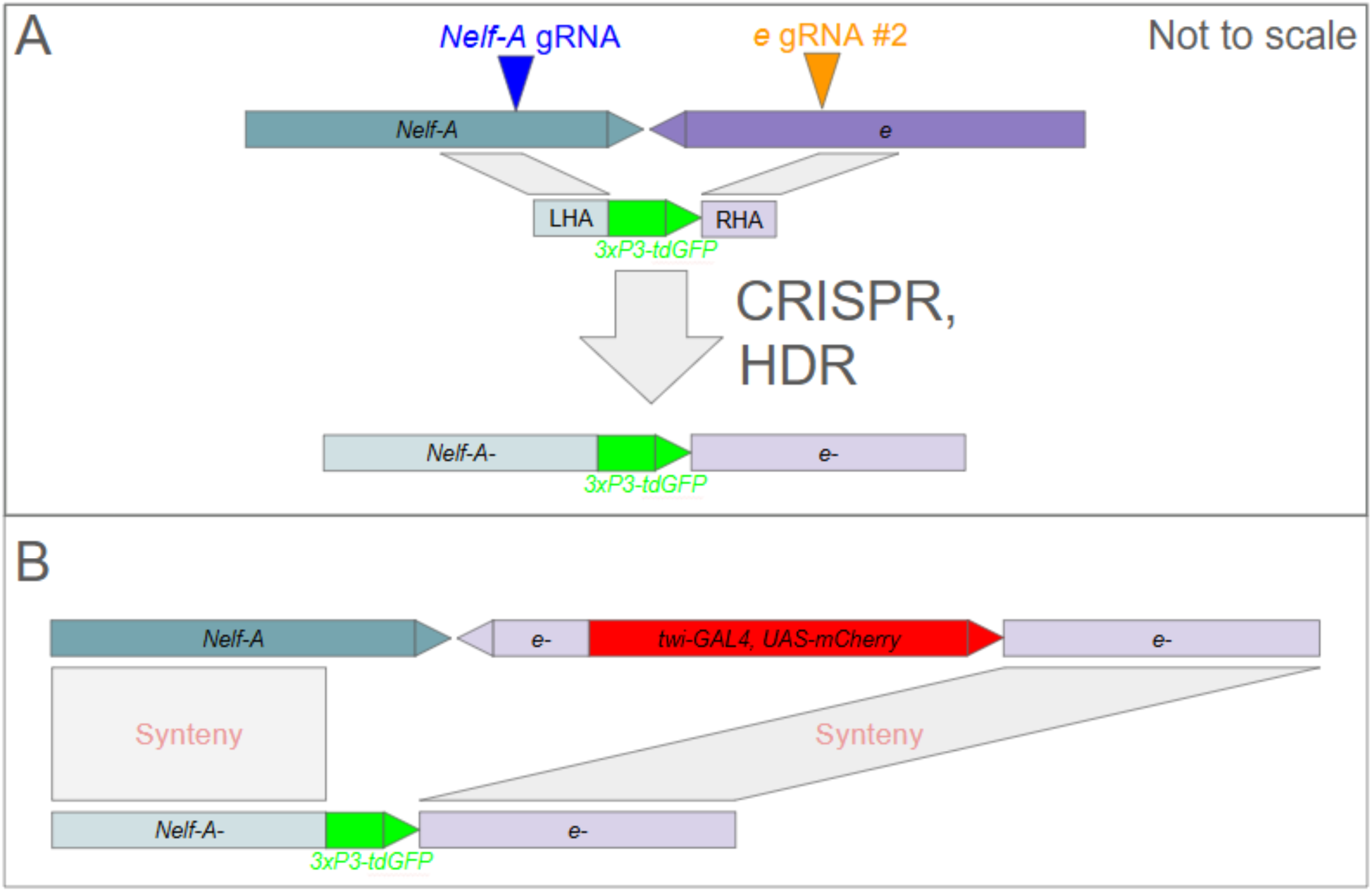
A) deletion of Nelf-A and replacement with 3xP3-tdGFP via CRISPR; B) visualization of the genotype of the SPECIES *twi-GAL4, UAS-mCherry/Nelf-A-, e-{3xP3-tdGFP}* stock at the locus where the transgenes are inserted. Because both aberrations are overlapping and homozygous lethal (with no opportunity for recombination to restore full viability to either allele), the alleles “balance” one another, circumventing the difficulty of maintaining the homozygous lethal *twi-GAL4, UAS-mCherry* transgene in the SPECIES background where conventional balancers cannot be introduced.

**Supplemental File 1:**

● **Supplemental File 1, Table 1**: Raw data underlying Table 1. Counts of embryos and larvae from red fluorescent (RFP+), green fluorescent (GFP+), and white light (Total, Larvae) images. Note that these categories are not exclusive (i.e. GFP+ embryos are a subset of RFP+ embryos, which in turn are a subset of the Total). The flies screened for drive were F1s between marker line males and GDL (or SD[72]) females.
● **Supplemental File 1, Table 2**: Sequences of gRNAs and homology arms used for CRISPR mutagenesis in this study.
● **Supplemental File 1, Table 3**: Camera settings used in image acquisition for embryo counting.
● **Supplemental File 1, Table 4**: Primers used to confirm proper integration of transgenes in Drosophila stocks created for this study.
● **Supplemental File 1, Table 5**: Sequences of plasmids injected as part of CRISPR- or phiC31-mediated mutagenesis.

